# Interaction between polygenic liability for schizophrenia and childhood adversity influences daily-life emotional dysregulation and psychosis proneness

**DOI:** 10.1101/778761

**Authors:** Lotta-Katrin Pries, Boris Klingenberg, Claudia Menne-Lothmann, Jeroen Decoster, Ruud van Winkel, Dina Collip, Philippe Delespaul, Marc De Hert, Catherine Derom, Evert Thiery, Nele Jacobs, Marieke Wichers, Ozan Cinar, Bochao D. Lin, Jurjen J. Luykx, Bart P. F. Rutten, Jim van Os, Sinan Guloksuz

**Affiliations:** Department of Psychiatry and Neuropsychology, School for Mental Health and Neuroscience, Maastricht University Medical Centre, Maastricht, the Netherlands; University Psychiatric Centre KU Leuven, Department of Neurosciences, KU Leuven, Belgium; University Psychiatric Centre Sint-Kamillus Bierbeek, Brothers of Charity, Belgium; Centre of Human Genetics, University Hospitals Leuven, KU Leuven, Belgium; Department of Obstetrics and Gynecology, Ghent University Hospitals, Ghent University, Belgium; Department of Neurology, Ghent University Hospital, Ghent University, Ghent, Belgium; Faculty of Psychology and Educational Sciences, Open University of the Netherlands, Heerlen, The Netherlands; University of Groningen, University Medical Center Groningen, Department of Psychiatry, Interdisciplinary Center Psychopathology and Emotion regulation (ICPE), The Netherlands; Department of Translational Neuroscience, UMC Utrecht Brain Center, University Medical Center Utrecht, Utrecht University, Utrecht, The Netherlands; Department of Psychiatry, UMC Utrecht Brain Center, University Medical Center Utrecht, Utrecht University, Utrecht, The Netherlands; GGNet Mental Health, Apeldoorn, the Netherlands, Medical Center Utrecht, Utrecht University, Utrecht, The Netherlands; King’s College London, King’s Health Partners, Department of Psychosis Studies, Institute of Psychiatry, London, United Kingdom; Department of Psychiatry, Yale School of Medicine, New Haven, CT

**Keywords:** Gene-environment interaction, ecological momentary assessment, psychosis, childhood adversity, stress, polygenic risk score

## Abstract

**Background:** The earliest stages of the pluripotent psychopathology on the pathway to psychotic disorders is represented by emotional dysregulation and subtle psychosis expression, which can be measured using the Ecological Momentary Assessment (EMA). However, it is not clear to what degree common genetic and environmental risk factors for psychosis contribute to variation in these early expressions of psychopathology.

**Methods:** In this largest ever EMA study of a general population twin cohort including 593 adolescents and young adults between the ages of 15 and 35 years, we tested whether polygenic risk score for schizophrenia (PRS-S) interacts with childhood adversity (the Childhood Trauma Questionnaire score) and daily-life stressors to influence momentary mental state domains (negative affect, positive affect, and subtle psychosis expression) and stress-sensitivity measures.

**Results:** Both childhood adversity and daily-life stressors were associated with increased negative affect, decreased positive affect, and increased subtle psychosis expression, while PRS-S was only associated with increased positive affect. No gene–environment correlation was detected. We have provided novel evidence for interaction effects between PRS-S and childhood adversity to influence momentary mental states [negative affect (b = 0.07, 95% CI 0.01 to 0.13, *P* = 0.013), positive affect (b = −0.05, 95% CI −0.10 to −0.00, *P* = 0.043), and subtle psychosis expression (b = 0.11, 95% CI 0.03 to 0.19, *P* = 0.007)] and stress-sensitivity measures.

**Conclusion:** Exposure to childhood adversities, particularly in individuals with high PRS-S, is pleiotropically associated with emotional dysregulation and psychosis proneness.

## Introduction

Converging evidence suggests that the genetic and nongenetic vulnerability contributing to the development of schizophrenia and related psychotic disorders is shared across a broad range of psychotic and non-psychotic clinical syndromes and expressed non-specifically in the affective, psychotic, and cognitive domains in the general population(1–5). Understanding the pleiotropic effects of risk factors associated with schizophrenia on the earliest stages of pluripotent psychopathology may therefore pave the way for gaining insight into the shared biological and mental processes underlying psychosis spectrum disorder (PSD). Contemporary concepts of mental disorders acknowledge psychopathology as a highly dynamic and time-varying complex system that can only be understood from its interconnected constituent parts. These concepts provide a useful theoretical framework to investigate how alterations of micro-level transdiagnostic mental states, varying from moment-to-moment, precede the transition to the more discrete clinical syndrome of PSD(6).

Studies of Ecological Momentary Assessment (EMA), designed to collect micro-level mental state variation, have consistently shown that disturbed emotional (affective dysregulation) and psychotic reactivity to daily-life stressors (aberrant salience attribution) are associated with psychosis expression in different populations at varying severity stages, the general population, clinical high-risk samples, siblings of patients with PSD, and cases(7–13). Further, in agreement with the diathesis-stress theory(14), EMA studies have provided evidence that genetic and environmental vulnerabilities are associated with alterations in emotional reactivity. Individuals who experienced childhood adversity (CA) showed heightened emotional and psychotic reactivity to daily-life stressors(15–19), increased persistence of momentary mental states(20); and the influence of CA on the reactivity to daily-life stressors were stronger in populations with increased proxy genetic risk (i.e. service users, clinical high-risk, or first episode psychosis compared to healthy controls)(15, 16).

Early hypothesis-driven candidate gene studies also provided some evidence for the role of gene-environment interaction (G×E) in affective and psychotic reactivity to daily-life stressors(21–23). While these studies can be considered the first steps in understanding the genetic correlates of daily stress-reactivity, they were undersized and by design too simplistic to capture the complex genetic architecture. The use of cumulative risk scores—polygenic risk score (PRS)—as a single molecular metric has significantly enhanced the power to detect G×E without compromising the validity of the results(24). We previously showed that the likelihood of schizophrenia is increased as a function of the interaction between PRS for schizophrenia (PRS-S) and childhood adversities as well as cannabis use(25).

A recent perspective article discusses how real-time measurement of cognitive and emotional processes via EMA, which eliminates retrospective recall bias, combined with modern polygenic approach may greatly advance our understanding of the role of G×E in psychopathology and mental wellbeing(26). PRS-based approaches for testing G×E represent a novel approach, and to the best of our knowledge, no EMA study has utilized PRS-S yet.

In this study, guided by the transdiagnostic Research Domain Criteria (RDoC) framework prioritizing shared dimensional psychological constructs cutting across diagnostic categories(27), we outlined a step-by-step analytical plan to test the contribution of G×E to altered emotional processes, previously associated with the earliest stages of PSD. Bringing together a unique sampling frame of a general population twin cohort of young adults and adolescents with rich EMA data, we aimed to investigate for the first time whether molecular genetic risk score for schizophrenia (PRS-S) interacts with early-life stressors (CA) and daily-life stressors (social, event, activity, and overall stress) to influence momentary mental state domains (negative affect, positive affect, and subtle psychosis expression), and whether PRS-S moderates the association between CA and stress-sensitivity measures.

## Methods

### Sample

Data were derived from the first wave of the TwinssCan, a general population twin cohort that started including adolescent and young adult (age range = 15-35 years) twins (n = 796), their siblings (n = 43), and parents (n = 363) from April 2010 to April 2014(28). The TwinssCan cohort comprises individuals fulfilling the inclusion criteria from the East Flanders Prospective Twin Survey(29), a prospective population-based, multi-birth registry positioned in Flanders, Belgium. Participants were excluded if they had a pervasive mental disorder as indicated by caregivers. All participants gave written informed consent and parent(s) signed an informed consent for participants below the age of 18 years. The local ethics committee approved the study (Commissie Medische Ethiek van de Universitaire ziekenhuizen KU Leuven, Nr. B32220107766). Sequential analysis based on sex, fetal membranes, umbilical cord blood groups, placental alkaline phosphatase, and DNA fingerprints was used to determine zygosity(29).

### Measures

#### Ecological Momentary Assessment (EMA)

EMA is a well-validated structured diary technique that assesses individual and contextual measures in the current moment, throughout the day(30–32). During the assessment period (6 consecutive days), participants used a digital device (PsyMate)(33) to electronically fill out a brief questionnaire assessing their emotions, thoughts, context and their appraisal of that context 10 times/day at an unpredictable moment (semi-random) in each of ten 90-minute time blocks between 7:30 and 22:30(33).

Conforming to previously described methods, the negative affect (NA)(15) and the positive affect (PA)(34) domains were the mean scores of items assessing emotional states. Subtle psychosis expression (PE) was the mean score of items concerning psychotic-like experiences(35). Daily-life stress domains were constructed as event(36), social(15), activity(15), and overall stress (average of event, social, and activity stress). For detailed description of EMA items, see **Table 1**.

**Table 1.** Description of EMA variables

|  |  |
| --- | --- |
| **Negative affect** | Mean score of 5 items (I feel anxious, lonely, down, insecure and irritated). Each item was rated on a 7-point Likert scale ranging from 1 (not at all) to 7 (very). |
| **Positive affect** | Mean score of 4 items (I feel cheerful, satisfied, relaxed, and globally feeling well). Each item was rated on a 7-point Likert scale ranging from 1 (not at all) to 7 (very). |
| **Subtle psychosis expression** | Mean score of 5 items (suspiciousness, being afraid of losing control, racing thoughts, pervasive thoughts, and difficulties to express thoughts). Each item was rated on a 7-point Likert scale ranging from 1 (not at all) to 7 (very). |
| **Event stress** | Participants were asked to rate the most important event since last entry on its pleasantness on a bipolar Likert scale from -3 (very unpleasant) to 3 (very pleasant). Consistent with previous studies(36), ratings from -3 to -1 were considered stressful and scores from 1 to 3 were recoded to 0. For simplification, the score was reversed and very unpleasant events therefore represented the highest score on a scale from 0-3. |
| **Social stress** | Participants were asked with whom they currently were (e.g., nobody or family). When participants reported to be alone, they were asked to answer the following items: I like to be alone (reversed); I would prefer to have company; and I feel safe alone (reversed). When participants reported to be in company, they were asked the following items: I would prefer to be alone; I find the people I am with pleasant (reversed); I feel safe (reversed); I feel I belong (reversed); and I feel judged. All items were scored on a Likert scale from 1 (not at all) to 7 (very), and the mean score of all items was used to estimate the social stress. |
| **Activity stress** | Participants were asked about the activity they participated in just before the beep (e.g., resting, watching TV, and smoking). The mean score of the following items [rated on a 7-point Likert scale from 1 (not at all) to 7 (very)] was used to calculate an activity stress score: I would prefer doing something else; This activity is difficult for me; and I can do this well (reversed). |
| **Overall stress** | Average of event-, social-, activity stress. |

EMA: Ecological momentary assessment

#### Childhood adversity

CA was assessed using the Childhood Trauma Questionnaire (CTQ)(37) that consists of 28 items rated on a 5-point Likert scale assessing five domains of maltreatment (emotional and physical neglect along with emotional, physical, and sexual abuse). **Supplementary Table 3** reports the frequencies of childhood adversity domains. Consistent with previous work(38) using this dataset, CA was defined as the mean score of all five domains.

### Genotyping, imputation, and PRS

Genotypes of the twins and their siblings were generated on two platforms: the Infinium CoreExome-24 and Infinium PsychArray-24 kits. Quality control (QC) procedures were performed using PLINK v1.9(39) in both datasets separately. Single nucleotide polymorphisms (SNPs) and participants with call rates below 95% and 98%, respectively, were removed. A strict SNP QC only for subsequent sample QC steps was then conducted. This involved a minor allele frequency (MAF) threshold > 10% and a Hardy-Weinberg equilibrium (HWE) *P*-value > 10^−5^, followed by linkage disequilibrium (LD)-based SNP pruning (R^2^< 0.5). This resulted in ~ 58K SNPs to assess sex errors (n=8), heterozygosity [F<5× the standard deviation (SD), n = 3], homozygosity (F > 5× SD), and relatedness by pairwise identity by descent (IBD) values (monozygotic: 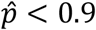, dizygotic and full siblings: 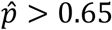 or 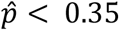, n = 5). The ancestry-informed principal components (PCs) analyses were conducted by EIGENSTRAT(40). The ethnic outliers of which the first 4 PCs diverged > 10× SD from Utah residents with Northern and Western European ancestry from the CEPH collection (CEU) and Toscani in Italia (TSI) samples (n = 5), and > 3× SD of the TwinssCan samples (n = 7) were excluded. After removing these subjects, a regular SNP QC was performed (SNP call rate > 98%, HWE p > 1e-06, MAF > 1%, and strand ambiguous SNPs and duplicate SNPs were removed).

The two QCed datasets were imputed on the Michigan server(41) using the HRC r1.1 2016 reference panel with European samples after phasing with Eagle v2.3. Post-imputation QC involved removing SNPs with imputation quality (R^2^) < 0.8, with a MAF < 0.01, SNPs that had a discordant MAF compared to the reference panel (MAF difference with HRC reference > 0.15), as well as strand ambiguous AT/CG SNPs and multi-allelic SNPs. The two chips were merged and an additional check for MAF > 0.01, HWE *P* > 1e-06 was executed, which resulted in 3,407,392 SNPs for 688 individuals.

PRS-S were calculated based on the meta-analysis results from the Psychiatric Genetics Consortium (PGC)-2 SZ and the CLOZUK sample (schizophrenia cases on clozapine from the UK)(42). Then insertions and deletions, ambiguous SNPs, SNPs with a MAF < 0.01, imputation quality R^2^< 0.9, SNPs located in complex-LD regions and long-range LD regions(43) were excluded. Overlapping SNPs between the schizophrenia GWAS (training), 1000 genomes (reference), and our dataset (target) were selected. These SNPs were clumped in two rounds using PLINK’s clump function (round 1: --clump-kb 250 --clump-r2 0.5; round 2: --clump-kb 5000 --clump-r2 0.2), resulting in 88,736 SNPs for PRS-S calculation. Odds ratios for autosomal SNPs reported in the schizophrenia summary statistics were log-converted into beta values. PRS-S were calculated using PLINK’s score function. Informed by the PGC analyses, PRS-S with cut-off *P* < 0.05 (including 21,901 SNPs) was used in the following analyses to achieve a balance between the number of false-positive and true-positive risk alleles(44). For details, see supplementary information.

### Statistical analyses

For the purpose of this analysis, parents were excluded. Only participants with complete data on the CTQ, age, sex, and PRS-S were included in the analyses. Conforming to previous studies(32), participants who completed less than 1/3 of the EMA questionnaires were excluded from the analysis (n = 52). One individual with a visibly extreme value of CA (> 7 SD from the mean) was excluded from analyses. The final sample included 593 participants: monozygotic (n = 180), dizygotic (n = 380) twin pairs, and their siblings (n = 33).

The data have a hierarchical structure. Multiple EMA observations (level 1) were clustered within subjects (level 2), who were part of twin pairs (level 3). Multilevel mixed-effects model is the recommended method to handle data including observations at more than one level in terms of unit of analysis by taking into account of the variability associated with each level of nesting(45–47). To handle this nested structure including familial relatedness, multilevel mixed-effects models were applied. In the current study, as typically observed in EMA studies, left-censoring (NA or PE) and right-censoring (PA) were present due to a greater amount of observations with a score of one (NA or PE) or seven (PA) on the outcome variables. In consideration of the skewness, multilevel mixed tobit regression(48) (censored regression) with an unstructured covariance matrix was performed using the Stata version 15.0(49) “METOBIT” command. The independent variables [PRS-S, CA, and daily-life stressors (overall, event, social, and activity stress)] were standardized and centered (min = 0, SD = 1).

First, we analyzed associations of CA and PRS-S, and their interaction, with EMA outcomes. Second, we tested associations of daily-life stressors, and their interaction with PRS-S, and EMA outcomes. Third, for sensitivity analyses, we constructed stress-sensitivity measures for use in G×E analyses. Consistent with previous work(50), separate models including the daily-life stressors as independent variables and EMA outcomes as dependent variables were estimated; fitted values of these models (substituting maximum likelihood estimates for fixed effects and empirical Bayes predictions for random effects) were stored as stress-sensitivity (e.g. NA-Event stress-sensitivity) scores. Eventually, we tested associations of CA and PRS-S, and their interaction, with normally distributed stress-sensitivity measures as dependent variables in multilevel linear regression models using the “MIXED” command. All models were controlled for *a priori* covariates (age and sex), while models including PRS-S were additionally adjusted for ancestry, using the first 2 genomic principal components (PCs). To adequately control for confounding(51), interaction models included these covariates not only as main effects but also covariate x environment and covariate x PRS-S interaction terms.

### Results

Sample characteristics are reported in **Table 2**. A correlation matrix of the three momentary mental state domains is provided in **Supplementary Table 4.**

**Table 2.** Sample characteristics

|  | Total sample (N = 593)<br>Mean (SD), EMA observations |
| --- | --- |
| Sex |  |
| Female | 362 (61%) |
| Male | 231 (39%) |
| Age | 17.60 (3.81) |
| Childhood adversity | 1.35 (0.31) |
| Negative affect | 1.78 (0.84), n = 23293<sup>a</sup> |
| Positive affect | 5.06 (1.06), n = 23265<sup>a</sup> |
| Subtle psychosis expression | 1.89 (0.99), n = 23272<sup>a</sup> |
| Overall stress | 1.89 (0.78), n = 23225<sup>a</sup> |
| Event stress | 0.34 (0.84), n = 22812<sup>a</sup> |
| Social stress | 2.33 (1.08), n = 23172<sup>a</sup> |
| Activity stress | 2.95 (1.43), n = 23192<sup>a</sup> |

<sup>a</sup> Number of observations, EMA: Ecological momentary assessment, PRS-S: Polygenic risk score for schizophrenia, SD: Standard deviation

### Main associations and interactions of CA and PRS-S on momentary mental state domains

CA was associated with increased NA, decreased PA, and increased PE, while PRS-S was only associated with increased PA **(Table 3)**. These results remained significant after controlling for daily-life stressors **(Supplementary Table 5**). No gene–environment correlation was present as PRS-S was not associated strongly or significantly with CA (b = − 0.01, 95% CI −0.03 to 0.02, *P* = 0.676).

**Table 3.** Associations and interaction effects of CA and PRS-S with momentary mental state domains

|  | Association with CA |  |  | Association with PRS-S<sup>a</sup> |  |  | Interaction between PRS-S and CA<sup>a</sup> |  |  |
| --- | --- | --- | --- | --- | --- | --- | --- | --- | --- |
|  | b | *P*-value | 95% CI | b | *P*-value | 95% CI | b | *P*-value | 95% CI |
| **Negative affect** | 0.13 | < 0.001* | 0.07 to 0.19 | -0.02 | 0.502 | -0.08 to 0.04 | 0.07 | 0.013* | 0.01 to 0.13 |
| **Positive affect** | -0.12 | < 0.001* | -0.17 to -0.06 | 0.08 | 0.003* | 0.03 to 0.14 | -0.05 | 0.043 | -0.10 to -0.00 |
| **Subtle psychosis expression** | 0.18 | < 0.001* | 0.10 to 0.26 | -0.03 | 0.547 | -0.11 to 0.06 | 0.11 | 0.007* | 0.03 to 0.19 |

All analyses were adjusted for age and sex, <sup>a</sup>also adjusted for 2 principal components, \*significant after controlling for family-wise type-I error using the Bonferroni method (0.05/3=0.0167). CA: Childhood adversity, CI: Confidence interval, PRS-S: Polygenic risk score for schizophrenia

PRS-S moderated the association of CA with all three momentary mental state domains, while only NA and PE reached the Bonferroni adjusted statistical significance level (**Table 3**). The interaction effects remained significant after controlling for daily-life stressors (**Supplementary Table 5**). As shown in **Figure 1**, visualizing the fitted interaction effects between PRS-S and CA on momentary mental state domains, the association between CA and mental state domains increased as a function of increased PRS-S (for scatter plots of raw data, see **Supplementary Figure 3).**

**Figure 1:**
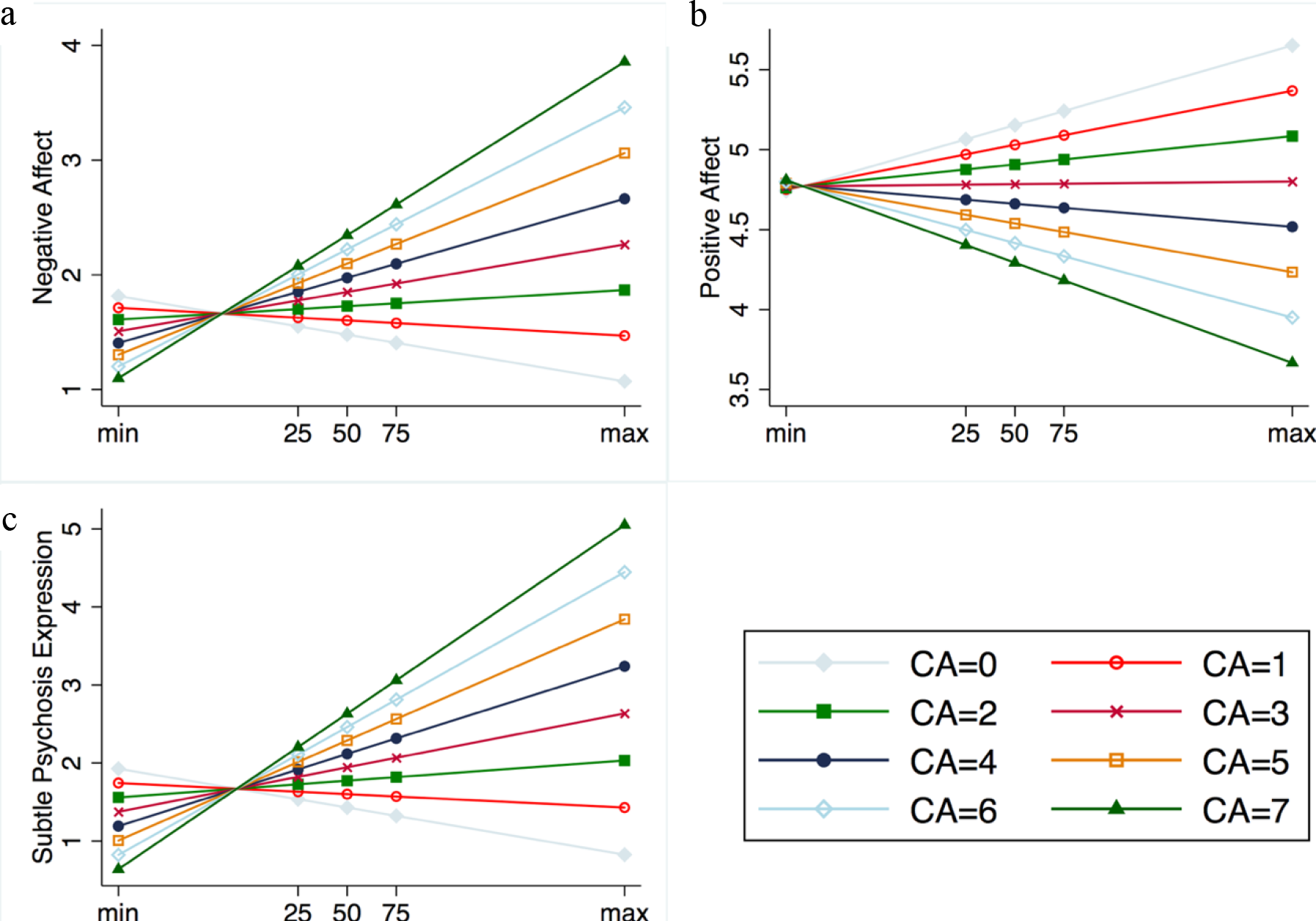
Interaction effect of CA and PRS-S on momentary mental state domains. Marginal effect plots based on multilevel tobit regression of the interaction between continuous polygenic risk score for schizophrenia (x-axis) and continuous childhood adversity score on continuous measures of negative affect (a), positive affect (b) and subtle psychosis expression (c), y-axis). For visualization purposes, margins at quartiles of PRS-S and standardized scores of CA from 0 to 7 were illustrated. CA: Childhood adversity, PRS-S: Polygenic risk score for schizophrenia [i.e. range: min (minimum), 25th percentile, 50th percentile, 75th percentile, and max (maximum)].

### Main associations and interactions of daily-life stressors and PRS-S on momentary mental state domains

The overall mean and each of the daily-life stressors were associated with increased NA, decreased PA, and increased PE **(Supplementary Table 6)**. No gene–environment correlation was present as PRS-S was not associated with any of the daily-life stressors (overall stress: b = −0.02, 95% CI −0.06 to 0.03, *P* = 0.491; event stress: b = 0.01, 95% CI −0.04 to 0.06, *P* = 0.672; social stress: b = −0.03, 95% CI −0.07 to 0.02, *P* = 0.267; activity stress: b = −0.01, 95% CI −0.05 to 0.03, *P* = 0.571). No evidence for significant interaction effects between daily-life stressors and PRS-S was found **(Supplementary Table 6)**.

### Main associations and interactions of CA and PRS-S on stress-sensitivity measures

CA was associated with increased stress-sensitivity measures, while PRS-S was only associated with increased PA stress-sensitivity. Evidence was found for significant gene-environment interaction. The associations between CA and stress-sensitivity measures was greater if individuals had higher PRS-S (**Table 4)**.

**Table 4.** Main associations and interaction effects of CA in the models of stress-sensitivity measures

|  | Association with CA |  |  | Association with PRS-S<sup>a</sup> |  |  | Interaction between PRS-S and CA<sup>a</sup> |  |  |
| --- | --- | --- | --- | --- | --- | --- | --- | --- | --- |
|  | b | *P*-value | 95% CI | b | *P*-value | 95% CI | b | *P*-value | 95% CI |
| NA-Overall | 0.12 | < 0.001<sup>*</sup> | 0.07 to 0.18 | -0.02 | 0.487 | -0.08 to 0.04 | 0.07 | 0.012<sup>*</sup> | 0.01 to 0.12 |
| NA-Event | 0.12 | < 0.001 | 0.07 to 0.18 | -0.02 | 0.493 | -0.08 to 0.04 | 0.07 | 0.013 | 0.01 to 0.12 |
| NA-Social | 0.12 | < 0.001 | 0.07 to 0.18 | -0.02 | 0.473 | -0.08 to 0.04 | 0.07 | 0.012 | 0.01 to 0.12 |
| NA-Activity | 0.12 | < 0.001 | 0.07 to 0.18 | -0.02 | 0.471 | -0.08 to 0.04 | 0.07 | 0.011 | 0.02 to 0.12 |
| PA-Overall | -0.11 | < 0.001<sup>*</sup> | -0.16 to -0.06 | 0.08 | 0.003<sup>*</sup> | 0.03 to 0.13 | -0.05 | 0.042 | -0.10 to -0.00 |
| PA-Event | -0.11 | < 0.001 | -0.16 to -0.06 | 0.08 | 0.003 | 0.03 to 0.13 | -0.05 | 0.042 | -0.10 to -0.00 |
| PA-Social | -0.11 | < 0.001 | -0.16 to -0.06 | 0.08 | 0.003 | 0.03 to 0.13 | -0.05 | 0.043 | -0.10 to -0.00 |
| PA-Activity | -0.11 | < 0.001 | -0.16 to -0.06 | 0.08 | 0.003 | 0.03 to 0.13 | -0.05 | 0.042 | -0.10 to -0.00 |
| PE-Overall | 0.18 | < 0.001<sup>*</sup> | 0.10 to 0.25 | -0.03 | 0.546 | -0.11 to 0.06 | 0.11 | 0.007<sup>*</sup> | 0.03 to 0.19 |
| PE-Event | 0.17 | < 0.001 | 0.10 to 0.25 | -0.03 | 0.546 | -0.11 to 0.06 | 0.11 | 0.006 | 0.03 to 0.19 |
| PE-Social | 0.18 | < 0.001 | 0.10 to 0.26 | -0.03 | 0.545 | -0.11 to 0.06 | 0.11 | 0.006 | 0.03 to 0.19 |
| PE-Activity | 0.18 | < 0.001 | 0.10 to 0.26 | -0.03 | 0.520 | -0.11 to 0.06 | 0.11 | 0.006 | 0.03 to 0.19 |

All analyses were adjusted for age and sex, <sup>a</sup>additionally adjusted for 2 principal components, <sup>\*</sup>significant after controlling for family-wise type-I error using the Bonferroni method (0.05/3=0.0167). CA: Childhood adversity, CI: Confidence interval, NA: Negative affect, PE: Subtle psychosis expression, PA: Positive affect, PRS-S: Polygenic risk score for schizophrenia

## Discussion

### Principal findings

In this first study testing PRS-S for an interaction with early and late stressors (childhood adversity and minor daily-life stressors) in association with dynamic pluripotent mental processes in the largest EMA dataset to date, evidence emerged for an interaction between PRS-S and childhood adversity to influence momentary mental states (negative affect, positive affect, and subtle psychosis expression) and stress-sensitivity measures.

### Stress exposure and emotional processes

In line with long-established findings from population-based datasets and samples of help-seeking adolescents and young adults(9, 15), we showed that minor daily-life stressors, regardless of the type of the stressor, were associated with all three domains of momentary mental states. More importantly, we provided further support for the shared vulnerability theory of mental disorders by demonstrating that CA was associated with NA, PA, and PE. These results echo recent findings from our line of research showing that CA is not exclusively associated with a specific mental disorder category, but rather with multidimensional psychopathology (cutting across diagnostic categories) in the general population, such as psychotic experiences, affective dysregulation, and negative symptoms(52–57). Therefore, it is plausible to conceptualize that the sensitivity to daily-life stressors is molded by previous exposure to significant life stressors as discussed in the models of diathesis-stress(14) and sensory processing sensitivity(58). Furthermore, the exposome, a dense network of environmental exposures(59, 60), may contribute to a person’s sensitivity to stress.

### Genetic vulnerability for schizophrenia moderates sensitivity to childhood adversity

PRS-S, as anticipated, had no significant predictive power for the EMA outcomes with the exception of positive affect-related items that were positively associated with PRS-S. The positive association between PRS-S and positive affect might seem counter-intuitive at first glance given that EMA studies have shown a decreased positive affect in patients with schizophrenia and PRS-S is associated with schizophrenia. However, emerging evidence suggests that the relation between PRS-S and symptom dimensions at the population level appears to be not following a simple logic. As this is the very first and the only EMA study investigating PRS-S, we could not make an exact comparison of our findings. Unfortunately, because most studies focus on psychopathology, it is also difficult to draw a parallel between our current results on positive affect and findings from studies investigating the association between PRS-S and symptom dimensions in healthy participants. These studies in healthy participants have shown inconsistent results(61–66). Some reported no association between PRS-S and several symptoms dimensions(63, 67), while others reported negative associations between polygenic risk and schizotypy(64, 67). In addition, in the rare instances where it has been examined, PRS-S appear to contribute to abilities required for a creative profession(68). Taken together, these findings in fact suggest a substantial fraction of pathoetiology may be explained by the influence of environment and the GxE. In agreement, our interaction analyses showed a consistent pattern. PRS-S moderated the influence of CA, but not the impact of minor daily-life stressors, for all momentary mental states, with negative affect and subtle psychosis expression reaching the Bonferroni adjusted statistical significance level. This interaction effect was similarly present in sensitivity to overall-stress and consistently observed for sensitivity to event, activity, and social stress.

The current results showing a difference between the degree of genetic moderation of two stressors (CA and daily-life stressors) underscores the importance of the type, timing, and extent of stressor in mental health impact. This is consistent with the neurodevelopmental hypothesis(69, 70) that postulates that exposure to early-life stressors in neurodevelopmentally sensitive periods are more likely to disturb the balance of important stress systems and lead to enduring emotional and behavioral problems in later life. These findings, combined with a recent meta-analysis showing that patients with PSD experience more negative emotion and less positive emotion in daily life(71), suggest that genetic and early adversities may have a permanent impact on mental well-being, resulting in a trait-like feature of person-specific alterations in emotional expression and psychosis proneness. An interesting question for future research spanning an extended period would be whether persistent low-threshold daily-life stressors may influence emotional reactivity toward mental ill-health in the long term or whether more serious life events are required to reach the threshold for clinical syndrome. Early candidate gene studies investigating the genetic moderation of mental health outcomes have generated mostly inconclusive findings due to methodological issues. COMT^Val158Met^ Val/Val carriers displayed increased paranoia in response to stress(22), while, in another study, Met/Met genotype was associated with increased PA in response to experiencing positive events(72). Momentary stress interacted with genetic variation in the brain-derived neurotrophic factor gene, Met carriers reporting higher paranoia scores than Val carriers(22). However, a recent study failed to replicate these findings, but showed an interaction between childhood trauma and *RGS4, FKBP5*, and *OXTR*, respectively(73). As PRS-based approaches for testing G×E have recently emerged, no comparable EMA study was available. However, in line with our study, research showed that higher PRS (consisting of 13 genes previously associated with vulnerability to environmental exposure) increased the influence of CA on stress-sensitization(74). Further, we recently showed evidence that the interaction between PRS-S and childhood adversity increases the likelihood of schizophrenia(25).

Given the influence of psychosocial stressors on immune processes and hypothalamic-pituitary axis modulation underlying the etiopathogenesis of PSD(75), future studies embracing biologically-informative target approaches may exploit the unique ability of EMA to capture dynamic fluctuation of mental states, and combine the granular information with multi-omics data (e.g., genome, proteome, and epigenome) to study candidate molecular mechanisms such as *FKBP5(76)* and extend previous EMA work investigating cortisol reactivity to daily-life stressors in relation to PSD(77).

### Pleiotropic influence of exposures and genetic vulnerability on psychopathology

Our findings agree with the literature showing that the influence of CA(78) and schizophrenia genetic liability(2, 79, 80) on mental health in the general population are pleiotropic and converge on shared psychological constructs and multidimensional psychopathology in the causal path to PSD. Considering the fact that mental health phenotypes (EMA outcomes in our study) are associated with each other at both dimensional and diagnostic levels and thereby violating the assumption of independence of pleiotropy, it is also plausible to argue that these disorders defined at the symptom level might be different expressions (phenotypic presentations) of a substantially shared pathoetiology with varying outcomes due to disease modifiers rather than distinct entities(81, 82).

A growing investment into transdiagnostic research of mental health will hopefully shed more light on this matter. Abundant evidence shows that the earliest psychopathological processes expressed before the prodrome of PSD are non-specific and include affective dysregulation, aberrant salience, and subtle cognitive disturbances(81). In this regard, EMA outcomes capturing subtle and transitory mental states, such as emotional reactivity and stress-sensitivity, are arguably more useful trans-diagnostic phenotypes than static questionnaire-based interval assessments to examine the contributions of environmental and genetic factors to variation in mental health at the community level(26). As recently proposed(83), multi-layered digital phenotyping via mobile devices may advance the RDoC work in the era of “Big Data” boosted by historic efforts of personalized medicine such as the National Institutes of Health initiative, the All of US research program.

### Limitations

The current study provided the first insights into the influence of genetic regulation of exposure to stressors on dynamic mental states by taking advantage of a unique population dataset with fine-grained phenotyping. However, several methodological considerations should be noted. First, although one of the strengths is that the sample comprises individuals at an age range when mental disorders often emerge, it is also possible that the association between PRS-S and stress may change as a function of aging and cumulative stressor load. Second, the retrospective collection of CA might be subject to recall and response biases; however, it is not intuitive how these would be differential with regard to EMA outcomes or PRS-S, or their interaction. Third, daily-life stressors might not only influence momentary mental states but might also be influenced by them. Fourth, EMA provides a unique opportunity to focus on moment-to-moment fluctuations of mental states; nevertheless, it may be more difficult to detect psychosis proneness than emotional reactivity in the general population.

### Conclusions

This observational study suggests that the exposure to childhood adversities, especially in individuals with high molecular genetic risk for schizophrenia, is associated with emotional dysregulation and psychosis proneness. Further pre-registered confirmatory research is required to validate these findings.

## Acknowledgments

The authors thank Jill Ielegems, Katrien Lyssens, Davinia Verhoeven, and Debora op’t Eijnde for data-collection. Further, the authors would like to acknowledge that the East Flanders Prospective Twin Survey (EFPTS) is partly supported by the Association for Scientific Research in Multiple Births and that the TwinssCan project is part of the European Community’s Seventh Framework Program under grant agreement No. HEALTH-F2-2009-241909 (Project EU-GEI).

