## Supplementary information for "Interaction between polygenic liability for schizophrenia and childhood adversity influences daily-life emotional dysregulation and psychosis proneness"

**Genotyping, quality control, imputation, and PRS**

Part 1 Target Genotype Data Processing

1. *Quality control for genotype data before imputation.*

TwinssCan data were genotyped on two chips: Infinium CoreExome-24 Kit (570,038 genotyped variants for 634 participants) and Infinium PsychArray-24 Kit (588,628 genotyped variants for 82 participants). These two datasets from different chips were quality controlled separately using PLINK v1.9(1)**.** Several pre-imputation quality control (QC) steps were applied to the datasets. In detail, single nucleotide polymorphisms (SNPs) and samples with call rates below 95% and 98%, respectively, were removed. A strict SNP QC only for subsequent sample QC steps was conducted. This involved a minor allele frequency (MAF) threshold > 10% and a Hardy-Weinberg equilibrium (HWE) $P$-value > $10^{-5}$, followed by linkage disequilibrium (LD) based SNP pruning ($R^2 < 0.5$). This resulted in ~58K SNPs to assess sex errors (n=8), heterozygosity [F<5x the standard deviation (SD), n=3], homozygosity (F>5x SD), and relatedness by pairwise identity by descent (IBD) values (monozygotic: $\hat{p} < 0.9$, dizygotic and full siblings: $\hat{p} > 0.65$ or $\hat{p} < 0.35$, n = 5). After removing failing samples, a regular SNP QC was performed (SNP call rate>98%, HWE p>1e-06, MAF>1%). Next, strand ambiguous SNPs and duplicate SNPs were removed. 270,976 variants and 610 individuals from chip 1, and 273,523 variants and 78 individuals from chip 2 passed these QC steps.

2. *Imputation on Michigan server.*

The two QC-ed datasets were converted into *.VCF files chunked by chromosome and imputed using the following settings: reference panel as HRC R1.1 2016; phasing as Eagle v2.3; population as European; model as QC & imputation. The imputation step resulted in 39,117,084 single nucleotide polymorphisms (SNPs). The general imputation quality is shown in **Supplementary Figure 1**.

*3. Quality control after imputation.*

The VCF files were firstly converted into PLINK best guess (hard call) genotypes by PLINK --vcf function. In the meantime, poor quality SNPs were excluded: multiallelic SNPs, SNPs with a minor allele frequency (MAF)<0.01 or INFO<0.8, and strand ambiguous AT/CG SNPs. The files were merged, remaining SNPs that passed QC from both arrays and another exclusion on MAF (<0.01) and Hardy Weinberger equilibrium (HWE) (<10e-6) in the whole dataset was performed. Finally, there were 3,407,392 SNPs and 695 individuals

*4. Principal components analyses (PCA).*

The principal components (PCs) analyses within TwinssScan samples, and TwinssScan samples along with Hapmap3 (as can be found online(2)) populations were conduct by EIGENSTRAT(3). A strict selection for SNPs which overlap with Hapmap3 SNPs list were conduct: 1. MAF>0.05, HWE>0.001; 2. Removal of 20 long LD regions (**Supplementary Table 1**); 3. LD pruned with an $R^2$ of 0.5; which resulting 95,466 best quality genotyped SNPs used to calculate genetic PCs.

PCs were firstly calculated with hapmap3 population to exclude European ethnic outliers: exceeding 10 times the standard deviation of Utah residents with Northern and Western European ancestry from the CEPH collection (CEU) and Toscani in Italia (TSI) populations for first 4 PCs. The first 2 PCs explained >80% of the total variance (see **Supplementary Table 2;** model 1). Five individuals are excluded from this dataset. Secondly, another PCA was conducted using the same SNP list (n=95,466) but calculated only in TwinssCan. The first 10 PCs explained >50% of the total variance (see **Supplementary Table 2;** model 2). Seven individuals were further considered as ethnic outliers by the first 4 PCs exceeding 3 times the standard deviation in the TwinssCan sample, and were excluded from this study. In the end, 688 individuals remained (see **Supplementary Figure 2**). After post-imputation quality control step, PCA was conducted using the same SNP list (n=95,466) within TwinssCan samples (n=688), and first two genetic PCs were further used to correct population stratification.

Part 2 Training schizophrenia GWAS summary statistic processing.

1. Training data quality control

PRS-S were calculated based on the summary statistics from the Psychiatric Genetics Consortium-2 SZ and the CLOZUK sample (schizophrenia cases from the UK)(4). GWAS summary statistic underwent multiple quality control steps and the odds ratios (ORs) were converted into beta values. The quality control criteria were following: 1) insertions and deletions, ambiguous SNPs; 2) SNPs with a MAF<0.01 and SNP with an imputation quality ($R^2$)<0.9 in training and target datasets; 3) Overlapping SNPs between the schizophrenia GWAS (training dataset), 1000 genomes (reference dataset), and our dataset (target dataset) were selected; 4) SNPs located in complex-LD regions(5) were excluded (**Supplementary Table 1**). These SNPs were clumped in two rounds using PLINK's clump function; round 1 with the default parameters (physical distance threshold 250kb and LD threshold ($R^2$) <0.5 (--clump-kb 250 --clump-r2 0.5); round 2 with a physical distance threshold of 5,000kb and LD threshold ($R^2$) < 0.2 (--clump-kb 5000 --clump-r2 0.2); resulting in 88,736 SNPs for PRS-S calculation.

2. Calculating PRS.

The beta-values, effective allele, and P-value were extracted from all summary statistics. The PRS was calculated using PLINK's score function for schizophrenia P-value threshold = 0.05.

**Supplementary Figure 1** Correlation of SNPs MAF from chip1 (Supplementary Figure 1 A) and chip 2 (Supplementary Figure 1 B) dataset with the reference MAF.

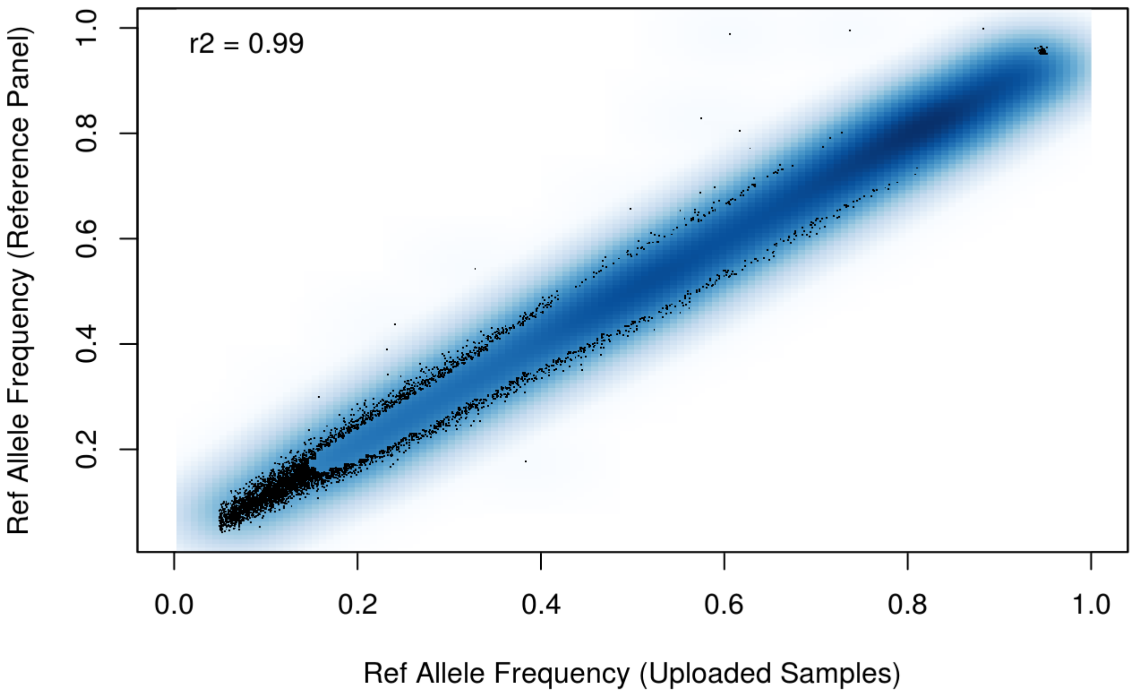

**Supplementary Figure 1 B** Correlation of SNPs MAF in chip2 dataset with the reference MAF.

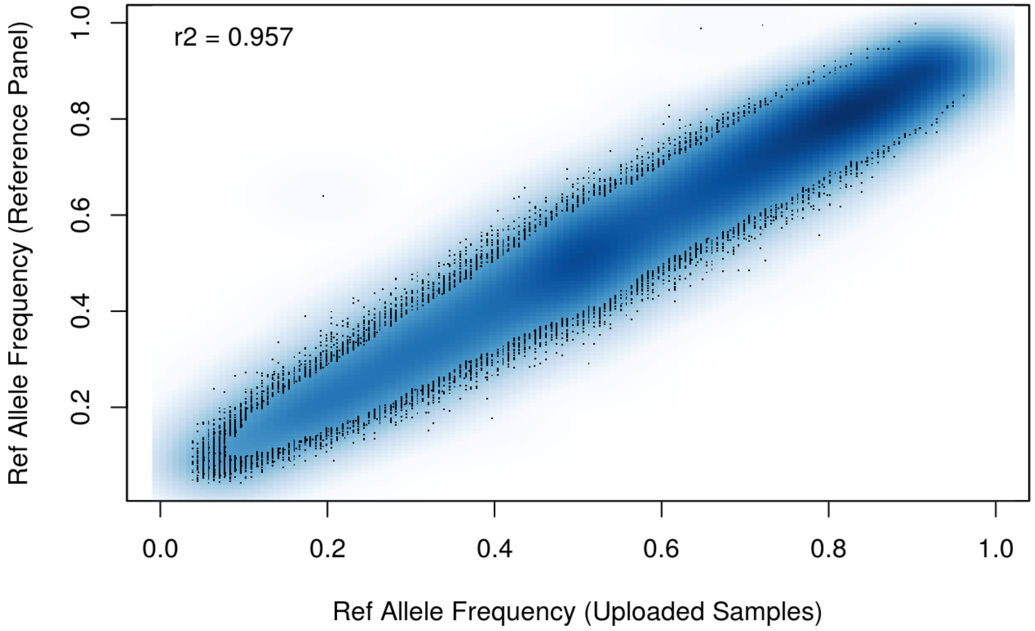

**Supplementary Figure 2** The first and second principal component of TwinssCan data (with identified ethnic outliers) along with hapmap3 populations.

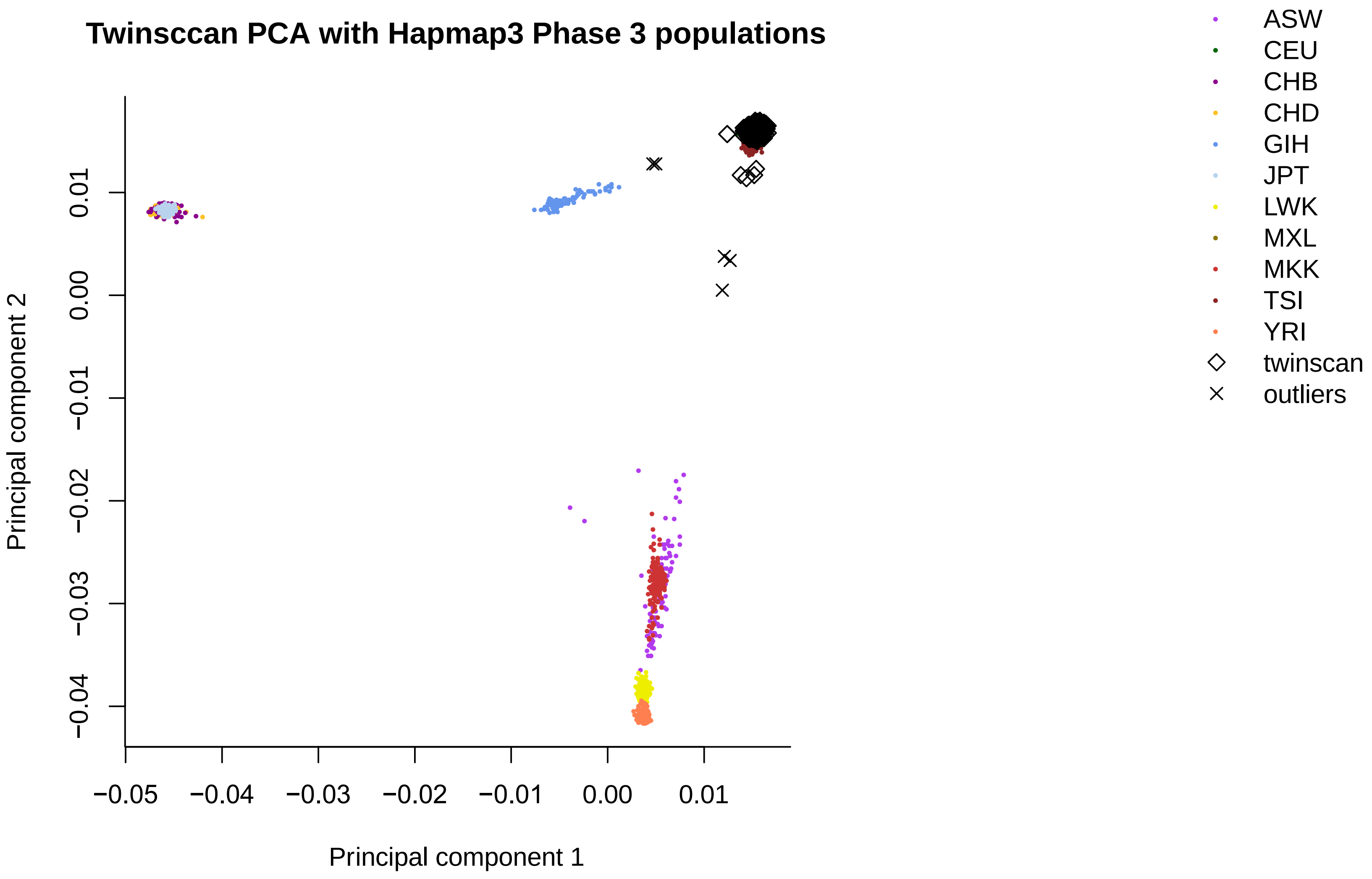

**Supplementary Table 1** 20 complex-LD regions and long-range LD regions which were exlcued from PRS analysis.

| Chromosome | Base pair position (start poiny to end point) |
|---|---|
| 1 | 48000000-52000000 |
| 2 | 86000000-100500000 |
| 2 | 183000000-190000000 |
| 3 | 47500000-50000000 |
| 3 | 83500000-87000000 |
| 5 | 44500000-50500000 |
| 5 | 129000000-132000000 |
| 6 | 25500000-33500000 |
| 6 | 57000000-64000000 |
| 6 | 140000000-142500000 |
| 7 | 55000000-66000000 |
| 8 | 8000000-12000000 |
| 8 | 43000000-50000000 |
| 8 | 112000000-115000000 |
| 10 | 37000000-43000000 |
| 11 | 87500000-90500000 |
| 12 | 33000000-40000000 |
| 20 | 32000000-34500000 |
| 8 | 8135000-12000000 |
| 17 | 40900000-45000000 |

**Supplementary Table 2** Eigenvalues and proportion variance explained for the first 20 PCs from PCA analyses.

| PCs | model 1: PCA with TwinssCan and hapmap3 population | | | model 2: PCA with TwinssCan cohort | | |
|---|---|---|---|---|---|---|
| | Eigenvalues | proportion variance | accumulated variance | Eigenvalues | proportion variance | accumulated variance |
| PC1 | 164.209 | 0.504 | 0.505 | 3.040 | 0.065 | 0.065 |
| PC2 | 101.007 | 0.310 | 0.815 | 2.748 | 0.058 | 0.123 |
| PC3 | 8.521 | 0.026 | 0.841 | 2.554 | 0.054 | 0.178 |
| PC4 | 7.795 | 0.023 | 0.865 | 2.482 | 0.053 | 0.231 |
| PC5 | 7.152 | 0.021 | 0.887 | 2.451 | 0.052 | 0.284 |
| PC6 | 2.773 | 0.008 | 0.896 | 2.427 | 0.051 | 0.336 |
| PC7 | 2.740 | 0.008 | 0.904 | 2.420 | 0.051 | 0.388 |
| PC8 | 2.692 | 0.008 | 0.912 | 2.364 | 0.050 | 0.438 |
| PC9 | 2.588 | 0.007 | 0.920 | 2.334 | 0.049 | 0.488 |
| PC10 | 2.490 | 0.007 | 0.928 | 2.210 | 0.047 | 0.536 |
| PC11 | 2.471 | 0.007 | 0.936 | 2.202 | 0.047 | 0.583 |
| PC12 | 2.430 | 0.007 | 0.943 | 2.188 | 0.046 | 0.630 |
| PC13 | 2.390 | 0.007 | 0.950 | 2.180 | 0.046 | 0.676 |
| PC14 | 2.351 | 0.007 | 0.958 | 2.174 | 0.046 | 0.723 |
| PC15 | 2.345 | 0.007 | 0.965 | 2.167 | 0.046 | 0.769 |
| PC16 | 2.312 | 0.007 | 0.972 | 2.158 | 0.046 | 0.815 |
| PC17 | 2.264 | 0.006 | 0.979 | 2.152 | 0.046 | 0.862 |
| PC18 | 2.259 | 0.006 | 0.986 | 2.149 | 0.046 | 0.908 |
| PC19 | 2.243 | 0.006 | 0.993 | 2.147 | 0.045 | 0.954 |
| PC20 | 2.224 | 0.006 | 1 | 2.145 | 0.045 | 1 |

**Supplementary Figure 3** Scatter plots of momentary mental state domains and PRS-S with lines of best fit at CA quartiles in raw data.

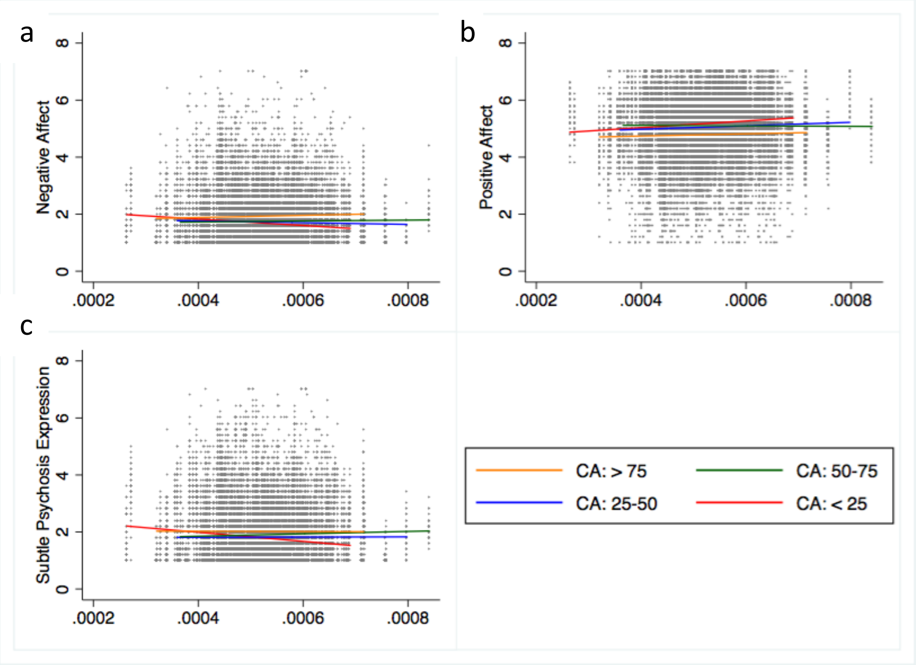

Interaction between polygenic risk for schizophrenia (x-axis) and childhood adversity on a) negative affect, b) positive affect and c) subtle psychosis expression (y-axis), CA: Childhood adversity (i.e. range: smaller than 25th percentile, between 25th percentile and 50th percentile, between 50th percentile and 75th percentile, and higher than 75th percentile), PRS-S: Polygenic risk score for schizophrenia.

**Supplementary Table 3** Childhood adversity subscales.

| | Emotional abuse (n = 593) | Physical abuse (N = 593) | Sexual Abuse (N = 593) | Emotional neglect (n = 593) | Physical neglect (N = 593) |
|---|---|---|---|---|---|
| None | 405 (68%) | 569 (96%) | 559 (94%) | 347 (59%) | 501 (84%) |
| Present | 188 (32%) | 24 (4%) | 34 (6%) | 246 (41%) | 92 (16%) |
| Low | 132 (70%) | 17 (71%) | 21 (62%) | 202 (82%) | 64 (70%) |
| Moderate | 35 (19%) | 5 (21%) | 9 (26%) | 32 (13%) | 20 (22%) |
| Severe | 21 (11%) | 2 (8%) | 4 (12%) | 12 (5%) | 8 (9%) |

Emotional abuse: None = 5-8, Low = 9-12, Moderate = 13-15, Severe ≥ 16; Physical abuse: None = 5-7, Low = 8-9, Moderate= 10-12, Severe ≥ 13; Sexual abuse: None = 5, Low = 6-7, Moderate = 8-12, Severe ≥ 13; Emotional neglect: None = 5-9, Low = 10-14, Moderate = 15-17, Severe ≥ 18; Physical neglect: None = 5-7, Low = 8-9, Moderate = 10-12, Severe ≥ 13

**Supplementary Table 4** Correlation matrix between momentary mental state domains.

| | Negative affect<br>rho, p-value | Positive affect<br>rho, p-value | Subtle psychosis expression<br>rho, p-value |
|---|---|---|---|
| Negative affect | 1 | -- | -- |
| Positive affect | -0.43, < 0.001 | 1 | -- |
| Subtle psychosis expression | 0.52, < 0.001 | -0.22, < 0.001 | 1 |

rho: Spearman's correlation

**Supplementary Table 5** Main and interaction effects of CA and PRS-S on momentary mental state domains adjusted for the individual daily-life stressors.

| | Association with CA | | | Association with PRS-S[a] | | | Interaction between PRS-S and CA[a] | | |
|---|---|---|---|---|---|---|---|---|---|
| | b | *P*-value | 95% CI | b | *P*-value | 95% CI | b | *P*-value | 95% CI |
| **Negative affect** | | | | | | | | | |
| Overall stress | 0.10 | < 0.001* | 0.05 to 0.15 | -0.01 | 0.642 | -0.06 to 0.04 | 0.07 | 0.003* | 0.02 to 0.12 |
| Event stress | 0.12 | < 0.001 | 0.06 to 0.18 | -0.02 | 0.492 | -0.08 to 0.04 | 0.08 | 0.006 | 0.02 to 0.13 |
| Social stress | 0.09 | 0.001 | 0.04 to 0.14 | -0.01 | 0.799 | -0.06 to 0.05 | 0.08 | 0.002 | 0.03 to 0.13 |
| Activity stress | 0.13 | < 0.001 | 0.07 to 0.18 | -0.02 | 0.525 | -0.07 to 0.04 | 0.07 | 0.009 | 0.02 to 0.12 |
| **Positive affect** | | | | | | | | | |
| Overall stress | -0.08 | < 0.001* | -0.13 to -0.04 | 0.07 | 0.006* | 0.02 to 0.12 | -0.05 | 0.037 | -0.09 to -0.00 |
| Event stress | -0.10 | < 0.001 | -0.16 to -0.05 | 0.08 | 0.004 | 0.03 to 0.14 | -0.06 | 0.019 | -0.11 to -0.01 |
| Social stress | -0.07 | 0.004 | -0.12 to -0.02 | 0.06 | 0.015 | 0.01 to 0.11 | -0.06 | 0.020 | -0.10 to -0.01 |
| Activity stress | -0.11 | < 0.001 | -0.16 to -0.07 | 0.07 | 0.006 | 0.02 to 0.12 | -0.05 | 0.050 | -0.09 to 0.00 |
| **Subtle Psychosis Expression** | | | | | | | | | |
| Overall stress | 0.15 | < 0.001* | 0.07 to 0.23 | -0.02 | 0.692 | -0.10 to 0.06 | 0.11 | 0.006* | 0.03 to 0.19 |
| Event stress | 0.17 | < 0.001 | 0.09 to 0.25 | -0.02 | 0.568 | -0.11 to 0.06 | 0.11 | 0.005 | 0.03 to 0.19 |
| Social stress | 0.15 | < 0.001 | 0.07 to 0.23 | -0.02 | 0.696 | -0.10 to 0.07 | 0.11 | 0.005 | 0.03 to 0.19 |
| Activity stress | 0.17 | < 0.001 | 0.09 to 0.25 | -0.02 | 0.660 | -0.10 to 0.06 | 0.11 | 0.006 | 0.03 to 0.19 |

Adjusted for age, sex, and daily-life stressors, [a]additionally adjusted for 2 principal components, *significant after controlling for family-wise type-I error using the Bonferroni method (0.05/3=0.0167). CA: Childhood adversity, CI: Confidence interval, PRS-S: Polygenic risk score for schizophrenia

**Supplementary Table 6** Main and interaction effects of daily-life stressors on momentary mental state domains.

| | Association with daily-life stressors | | | Interaction between PRS-S and daily-life stressors[a] | | |
|---|---|---|---|---|---|---|
| | b | *P*-value | 95% CI | b | *P*-value | 95% CI |
| **Negative affect** | | | | | | |
| Overall stress | 0.33 | < 0.001* | 0.31 to 0.35 | -0.00 | 0.851 | -0.02 to 0.02 |
| Event stress | 0.17 | < 0.001 | 0.15 to 0.19 | -0.02 | 0.086 | -0.04 to 0.00 |
| Social stress | 0.27 | < 0.001 | 0.25 to 0.29 | -0.00 | 0.809 | -0.02 to 0.02 |
| Activity stress | 0.24 | < 0.001 | 0.22 to 0.26 | -0.01 | 0.522 | -0.03 to 0.01 |
| **Positive affect** | | | | | | |
| Overall stress | -0.39 | < 0.001* | -0.41 to -0.37 | 0.02 | 0.145 | -0.01 to 0.04 |
| Event stress | -0.21 | < 0.001 | -0.23 to -0.19 | 0.02 | 0.172 | -0.01 to 0.04 |
| Social stress | -0.30 | < 0.001 | -0.32 to -0.28 | 0.01 | 0.220 | -0.01 to 0.03 |
| Activity stress | -0.29 | < 0.001 | -0.31 to -0.27 | 0.01 | 0.148 | -0.01 to 0.03 |
| **Subtle psychosis expression** | | | | | | |
| Overall stress | 0.21 | < 0.001* | 0.19 to 0.23 | -0.01 | 0.464 | -0.03 to 0.01 |
| Event stress | 0.10 | < 0.001 | 0.08 to 0.12 | -0.01 | 0.216 | -0.03 to 0.01 |
| Social stress | 0.17 | < 0.001 | 0.15 to 0.19 | -0.00 | 0.676 | -0.03 to 0.02 |
| Activity stress | 0.16 | < 0.001 | 0.14 to 0.18 | -0.01 | 0.388 | -0.03 to 0.01 |

Adjusted for age, sex, and daily-life stressors, [a]additionally adjusted for 2 principal components, [*]significant after controlling for family-wise type-I error using the Bonferroni method (0.05/3=0.0167).CA: Childhood adversity, CI: Confidence interval, PRS-S: Polygenic risk score for schizophrenia
